# Auditory Hair Cells and Spiral Ganglion Neurons Regenerate Synapses with Refined Release Properties In Vitro

**DOI:** 10.1101/2023.10.05.561095

**Authors:** Philippe F.Y. Vincent, Eric D. Young, Albert S.B. Edge, Elisabeth Glowatzki

**Affiliations:** The Center for Hearing and Balance; Department of Otolaryngology Head and Neck Surgery; Department of Neuroscience; Department of Biomedical Engineering, The Johns Hopkins School of Medicine, Baltimore, Maryland; Department of Otolaryngology, Harvard Medical School; Eaton-Peabody Laboratory, Massachusetts Eye and Ear, Boston, Massachusetts, USA; Program in Speech and Hearing Bioscience and Technology, Harvard Medical School, Boston, Massachusetts, USA; Harvard Stem Cell Institute, Cambridge, Massachusetts, USA

## Abstract

Ribbon synapses between inner hair cells (IHCs) and type I spiral ganglion neurons (SGNs) in the inner ear are damaged by noise trauma and with aging, causing ‘synaptopathy’ and hearing loss. Co-cultures of neonatal denervated organs of Corti and newly introduced SGNs have been developed to find strategies for improving IHC synapse regeneration, but evidence of the physiological normality of regenerated synapses is missing. This study utilizes IHC optogenetic stimulation and SGN recordings, showing that newly formed IHC synapses are indeed functional, exhibiting glutamatergic excitatory postsynaptic currents. When older organs of Corti were plated, synaptic activity probed by deconvolution, showed more mature release properties, closer to the highly specialized mode of IHC synaptic transmission that is crucial for coding the sound signal. This newly developed functional assessment of regenerated IHC synapses provides a powerful tool for testing approaches to improve synapse regeneration.

## INTRODUCTION

Inner ear ribbon synapses between inner hair cells (IHCs) and type I auditory nerve fibers (here referred to as spiral ganglion neurons (SGNs)) can be damaged during noise exposure or in the process of aging, this resulting in ‘synaptopathy’ that can contribute to hearing loss^1,2^. Noise exposure triggers massive glutamate release from IHCs, causing excitotoxicity via postsynaptic glutamate receptors on SGN endings. Several studies have investigated whether ribbon synapses do regenerate post noise-trauma and found that synapse regeneration does occur in several mammalian species^3–6^. In response to damaging noise exposure, SGNs retract from IHCs and can grow back toward IHCs to reform synapses. This regenerative process seems to be highly dependent on the level of sound exposure and can sometimes lead to poor regeneration after noise trauma.

Several studies have used *in vitro* cell-culture models of the cochlea to create new synaptic contacts and study mechanism that might facilitate synapse regeneration^7,8,9^. In one such model, early postnatal cochlear tissue was mechanically denervated and SGNs^10,11^ or stem cell-derived neurons^12,13^ were placed in co-culture. In such conditions, newly formed contacts between IHCs and newly added SGNs were confirmed by immunolabeling, showing the expression of both pre- and postsynaptic markers. Other studies demonstrated that neuronal progenitors obtained from embryonic stem cells promote ribbon synapse formation, both *in vitro* and *in vivo*^14,15^. Engraftment of embryonic stem-cell-derived otic progenitors into the cochlear nerve trunk of an adult gerbil model of deafness showed fiber growth to the IHCs^14^ and a partial restoration of the auditory hearing threshold^16^, as measured by auditory brainstem responses (ABRs). Several *in vivo* studies have further demonstrated protection of auditory synapses in a mouse model of noise-induced synaptopathy by genetic^17^ or viral^18^ delivery of NT3, the neurotrophin receptor agonist amitriptyline^19^ or by bisphosphonate^20^ before or within a short time-window after noise exposure. An antibody to Repulsive Guidance Molecule A (RGMa) was effective in restoring the synapse when administered onto the round window membrane one week after noise exposure^21^. So far, these studies have only relied on immunolabeling of pre-and postsynaptic markers for identifying newly formed IHC/SGN synapses as well as on ABR measurements. However, a direct functional measurement of the properties of such newly formed synaptic contacts has not been performed. To properly encode the sound signal, ribbon synapses must display fast, reliable and indefatigable synaptic transmission^22^. This is achieved by a complex release machinery^23^ associated with non-conventional postsynaptic properties that produce fast and unusually large EPSCs with various waveforms (mono- and multiphasic)^24–26^.

The goal of this study was to assess the physiological properties of each individual regenerated synaptic contact formed in co-cultures of postnatal mouse denervated organs of Corti and added SGNs. Hair cells were stimulated optogenetically and added SGNs were visualized by expression of fluorescent reporters, so their somata could be targeted for patch clamp recordings. Indeed, newly formed synapses were found to be functional, as light stimulation of hair cells activated glutamatergic synaptic currents in SGNs. Denervated organs of Corti were plated at different ages for co-culture to test if their age affects properties of synaptic transmission. Deconvolution of EPSC waveforms provided a powerful tool to probe for the mode of transmitter release and showed that, when older organs of Corti were plated, synaptic transmission in regenerated synapses showed more mature properties, closer to the specialized mode of native IHC synaptic transmission.

## MATERIALS AND METHODS

### Animals

All experiments were performed in accordance with protocols approved by the Johns Hopkins University Animal Care and Use Committee. Animals of either sex were used in the experiments indiscriminately. All mouse lines used is this study were maintained on a C57BL/6J background. Cochlear tissue with optogenetically competent auditory hair cells was obtained by crossing Growth Factor independent 1 Cre mice (*Gfi1^Cre/+^*; a gift from Dr. Lin Gan and Dr. Jian Zuo, New York^27^) with homozygous floxed Ai32 mice (*Ai32^fl/fl^*; Channelrhodopsin-2; ChR2; Jackson laboratory, #012569). Cochlear tissue with fluorescent SGNs was obtained from different mouse lines: 1) Microtubule-Associated Protein Tau driving the Green Fluorescent Protein expression (*Mapt-GFP^+/+^*, Jackson laboratory, #029219). MAPT is a protein involved in microtubule assembly and stability in neurons^28^; 2) the Basic Helix-Loop-Helix Family Member BHLHB5 Cre mice (*Bhlhb5^Cre/+^*^29,30^) crossed with homozygous floxed Ai9 mice (*Ai9^fl/fl^*; Td-Tomato, Jackson Laboratory #007905) and 3) Bhlhb5Cre/+ mice crossed with homozygous floxed GCaMP6f mice (*GCaMP6_f_^fl/fl^*; Jackson laboratory, #028865). BHLHB5 is a transcription factor expressed in the central nervous system and in sensory nerve fibers^31^.GCaMP6f is a genetically encoded Ca^2+^ indicator where GFP is coupled to the Ca^2+^ binding protein calmodulin. GCaMP6f displayed a basal GFP fluorescence that could be observed by epifluorescence with a 470 nm LED. Therefore, in this study, GCaMP6f was used as a GFP-like fluorescent reporter and not as a Ca^2+^ indicator due to cross interaction with the ChR2 excitation wavelength also at 470 nm.

### Preparation of denervated organs of Corti for culture

To prepare cultured cochlear tissue including inner and outer hair cells, without remaining connections to spiral ganglion neurons, a technique of culturing cochlear ‘micro-isolates’ first reported by Flores-Otero et al (2007)^32^ and later by Tong et al (2013)^10^ was adapted; here called ‘denervated organs of Corti’. At P3-P5, P7-P8 or P10-11, mice were euthanized, and cochlear tissue was extracted from each temporal bone of *Gfi1^Cre/+^; Ai32^fl/+^* or *Gfi1^+/+^; Ai32^fl/+^* mice, then transferred to a petri dish containing cold Hank’s Balanced Salt Solution (HBSS, Thermofisher, #14025-092). The organ of Corti, containing all IHC and OHC rows along with their surrounding supporting cell tissue, was physically ‘micro-isolated’ from the connecting afferent SGN fibers and spiral ganglion which includes the SGN somata, with a blade, close to the base of the IHCs. Denervated organs of Corti were obtained from the mid turn of the organ or Corti so that results could be compared between the different sessions. Denervated organs of Corti were then transferred onto coverslips that had been coated with 50 µg/ml of Laminin (Corning, #354232) and poly-L-Ornithine (0.01%, Sigma Aldrich, #P4957) in a 4-well petri dish (Greiner Bio One, #627170) and were cultured in DMEM/F12 – Glutamax (GIBCO, #10565-018) supplemented with N2 (GIBCO, #17502-048), B27 (GIBCO, #17504-044) and 10% Fetal Bovine Serum (FBS, GIBCO, #A3160401) and maintained in a humidified incubator with 5% CO2 at 37ᵒC for 4 hours. In a single session, typically 6 littermate mice were sacrificed, and 12 micro-isolates were plated onto 12 coverslips. For this study, 7 such sessions were performed with P_3-5_ cochlear tissue, 2 sessions with P_7-8_ cochlear tissue and 2 sessions with P_10-11_ cochlear tissue.

### Isolation of primary auditory neurons

SGNs were isolated by using a modified version of the previously described^12^. For fluorescently marked SGNs, either *Mapt-GFP^+/+^*, *Bhlhb5^Cre/+^; Ai9^fl/+^* or *Bhlhb5^Cre/+^; GCaMP6f ^fl/+^*mice were used (see Animals section). The cochlea was extracted from the temporal bone and transferred to a petri dish containing HBSS. Spiral ganglion tissue was isolated from six to eight P0-P2 mouse cochleas and pooled in a single tube. Briefly, the organ of Corti was removed from the basal to the apical turn, leaving the SGNs embedded in the soft modiolar connective tissue of the spiral ganglion, that was removed as much as possible before digestion. Spiral ganglion tissues were digested with 500 µl of Trypsin-EDTA 0.25% (GIBCO; #25200-056) and 500 µl of Collagenase-I 0.2% (GIBCO, #17018-029) during 20 min at 37°C. The digestion was stopped by adding 100 µl of FBS. The supernatant was carefully removed and replaced with fresh DMEM/F12 – Glutamax medium supplemented with N2, B27, 50ng/ml of Neurotrophin-3 (NT3, Novus biologicals; #NBP1 99227), 50ng/ml of Brain Derived Neurotrophic Factor (BDNF, Novus biologicals; #NBP1 99674) and 10% FBS. The mitotic inhibitor Cytarabine (AraC, 5µM; Sigma Aldrich, #1162002) was added to block glial cell proliferation and improve the SGN culture^33^. The antibody anti-Repulsive Guidance Molecule-A (RGMa, 10µg/ml; Thermofisher, #MA5-23977) was added, as it has been shown to promote synapse formation by blocking a repulsive guidance pathway^11^. Spiral ganglion tissues were then mechanically triturated by pipetting up and down several times. Using a cell counting chamber, this protocol allowed to isolate about 400.000 SGNs/ml from 8 cochleas. Then, about 30.000 SGNs were added to each well containing one coverslip with one micro-isolated hair cell tissue. The co-cultures were left for 10 to 12 days in the incubator until tested for newly formed synapses. During this time, culture medium was exchanged every 1 to 2 days.

### Electrophysiological recordings

At 10 to 12 days *in vitro* (DIV10 to DIV12), coverslips with co-cultures were placed in a recording chamber filled with an extracellular solution compatible with NMDA receptor activation (in mM): 144 NaCl, 2.5 KCl, 1.3 CaCl_2_, 0.7 NaH_2_PO_4_, 5.6 D-Glucose, 10 HEPES and 0.03 D-serine pH 7.4 (NaOH), 300 mOsm. Co-cultures were observed with a 40x water immersion objective (CFI60 Apo 40X W NIR, NA = 0.8, W.D = 3.5mm) attached to an upright Nikon A1R-MP microscope.

All patch clamp experiments were performed at room temperature using a double EPC10 HEKA amplifier controlled by the software Patchmaster (HEKA Elektronik). Patch pipettes, both for hair cell and SGN soma recordings, were pulled using a Flaming/Brown micropipette P-1000 puller (Sutter Instrument) and fire polished with a Micro forge MF-900 (Narishige) to achieve a resistance between 4-7 MΩ.

### Whole cell patch clamp recording from ChR2^+^ IHCs

Patch pipettes were filled with an intracellular solution (in mM): 145 KCl, 0.1 CaCl_2_, 1 MgCl_2_, 5 HEPES, 1 K-EGTA, 2.5 Na_2_-ATP, pH 7.4 (KOH), 300 mOsm. The holding membrane potential of IHCs was set at −74 mV (corrected for a liquid junction potential of 4 mV). IHCs of *Gfi1^Cre/+^; Ai32^fl/+^*mice were depolarized by activating channelrhodopsin-2 with a 470 nm blue LED (470/24 nm; 196mW) delivered by a Spectra X light engine (Lumencor Inc, Beaverton, OR) connected to the epifluorescent port of the Nikon A1R-MP microscope. The 470 nm light irradiance delivered through the 40x objective was estimated at 9mW/mm^2^ using a DET10A optical power meter (Thorlabs Newton, NJ).

### Recordings from SGN somata

Patch pipettes were filled with an intracellular solution containing the following (in mM): 110 K-MeSO_3_, 20 KCl, 0.1 CaCl_2_, 5 K-EGTA, 5 HEPES, 5 Na_2_-phosphocreatine, 5 Mg_2_-ATP, 0.3 Na_2_-GTP, pH 7.4 (KOH), 300 mOsm. Tetrodotoxin citrate (1-5 µM, #1069), (*RS*)-3-(2-Carboxypiperazin-4-yl)-propyl-1-phosphonic acid ((Rs)-CPP; 20 µM; #0173), an NMDA receptor blocker; 2,3-Dioxo-6-nitro-1,2,3,4-tetrahydrobenzo[*f*]quinoxaline-7-sulfonamide (NBQX; 20 µM; #1044), a glutamate −AMPA/Kainate receptor blocker and CP465-022 (1 µM; #2932), a selective AMPA receptor blocker, were purchased from R&D Systems. While TTX was added in the general perfusion, GluR blockers were locally perfused.

The SGN somata to record from, were selected by visualizing the hair cell rows and finding SGN fibers with fluorescing terminals close to the hair cell rows. From such fluorescing SGN endings, fibers were traced back to their somata, where patch recordings were made. SGN somata were randomly found from tens to hundreds of microns away from the hair cell-containing denervated organ of Corti tissue. For voltage clamp intracellular recordings, the holding potential of SGNs was set at −79 mV (corrected for a liquid junction potential of 9 mV). Recordings were included in the analysis if the holding current was < −200 pA at a holding potential of –79 mV, and if the series resistance (Rs) was < 20 MΩ. Recordings were not corrected for Rs.

Excitatory postsynaptic currents (EPSCs) were elicited by stimulating hair cells, either by optogenetic hair cell stimulation or by a local application of extracellular solution containing 40 mM K^+^, (sodium was reduced as potassium was increased). 40 mM K^+^-induced stimulation was used when hair cells were ChR2 negative. For some recordings, the absence of synaptic events in response to the optogenetic stimulation was confirmed by application of 40 mM K^+^.

### EPSC analysis

EPSC analysis was mostly performed as described previously^34^. After the baseline holding current was zeroed out, EPSCs were detected using a threshold at a manually-determined level (typically −10 to −20 pA), set by visual examination of data. Compared to EPSCs analyzed in the previous study, recordings obtained here produced EPSCs with smaller amplitudes on average, so it was not possible to find a threshold such that all or most EPSCs would exceed the threshold, while little or no noise did so. The threshold was set to eliminate noise as much as possible. Thus, a significant (and unknown) number of small EPSCs are not included in the data analyzed. A MATLAB routine provided EPSC amplitude, area, 10-90% rise times and 90-10% decay times for ‘fast’ EPSCs, both from recordings with purely fast and mixed (fast and slow) EPSCs. Only EPSCs from fast-type recordings were analyzed using deconvolution^34^ in which an

EPSC is expressed as the sum of one or more kernels. The kernel is the average of ‘monophasic’ EPSCs (defined as having one peak and a mono-exponential decay) from the individual recording, normalized to magnitude 1. An optimum summation of kernels to fit each EPSC is found by varying the amplitude and time delay of each kernel making up the sum. The optimization is done using the lasso method^35^, which uses the smallest number of kernels in the sum that produces a good fit to the EPSC waveform (least squares). The fitting process was done with the lasso routine in MATLAB. The assumption is that each kernel represents the release of a bolus of neurotransmitter. This analysis provides an estimate of the number of neurotransmitter releases in an EPSC and their arrangement in time and amplitude. Examples of fits determined with this method are shown in **Fig. 5A2-A5**. The green pulse trains in **Fig. 5A2-A5** show the amplitudes and times of occurrence of the kernels for four example EPSCs. The numbers show the number of kernels needed in each case. If multiple kernels are needed for fitting the EPSC, it is called multiphasic.

### Immunolabeling

P4-5 *Gfi1^+/+^; Ai32^fl/+^* isolated hair cells were co-cultured with wild-type P1 SGNs until DIV12. Co-cultures were fixed with 100% methanol for 10 min at −20°C. Next, coverslips were washed with 1X Phosphate Bovin Serum (PBS) and incubated with the blocking buffer containing 15% goat serum during 1h30 at room temperature. Tissues were stained to visualize presynaptic ribbons (Mouse IgG_1_ anti-CtBP2, 1:200; BD science: #612044; RRID: AB_399431), hair cells (Rabbit anti-Myosin VI, 1:200; Sigma #M5187; RRID: AB_260563), afferent nerve fibers (Chicken anti-NF200, 1:200; Millipore: #AB5539; RRID: AB_11212161), and the postsynaptic AMPA receptors (Mouse IgG_2B_ anti PAN-GluA1-4, 1:500; Millipore-Sigma: #MABN832). Coverslips were incubated with primary antibodies overnight at 4°C. Next, coverslips were thoroughly washed with 1X PBS before adding secondary antibodies (1:1000) for 2h at room temperature. The cocktail of secondary antibodies contained Goat anti-IgG1 Dylight 405 (Jackson Immuno Research: #115-477-185; RRID: AB_2632529), Goat anti-Rabbit Alexa 488 (Invitrogen: #A11008; RRID: AB_143165), Goat anti-Chicken CF405L (Biotium, custom made) and Goat anti-IgG_2b_ Alexa 647 (Invitrogen: #A21242; RRID: AB_2535811).

### Statistics

Statistical tests were performed with GraphPad Prism 9.2 (GraphPad Software, Inc). Data distribution was first tested for normality with a Shapiro-Wilk test. Parametric and non-parametric tests were then selected accordingly. When two populations were compared a student-t-test or a Mann-Whitney test was used. To compare more than two populations, a one-way ANOVA with a Tuckey post-hoc test or a Kruskal-Wallis with a Dunn’s post-hoc test was used. The limit of significance was set at p <0.05.

## RESULTS

### Co-cultures of denervated organs of Corti and isolated SGNs

To create regenerated synapses between hair cells and SGNs after synapse loss, co-cultures of denervated organs of Corti and isolated SGNs were established so that they allowed for new synaptic connections to form (modified from^10, 32^; **Fig. 1A**). Organs of Corti were denervated by cutting along the inner spiral bundle, where SGN endings contact the IHCs (**Fig.1A**, dotted red line). Denervated mid-turn organs of Corti were isolated from *Gfi1^Cre/+^; Ai32^fl/+^* and *Gfi1^+/+^; Ai32^fl/+^* (Channelrhodopsin-2; ChR2) mice, at postnatal days (P)3-5. In Gfi1^Cre^-positive organs of Corti, inner and outer hair cells express Channelrhodopsin-2 **(Fig. 1B),** allowing for hair cell transmitter release to be triggered by optogenetic stimulation. Additionally, in Gfi1^Cre^-negative organs of Corti, transmitter release was induced by local perfusion of extracellular solution with 40 mM K^+^ ^24^.

**Figure 1.**
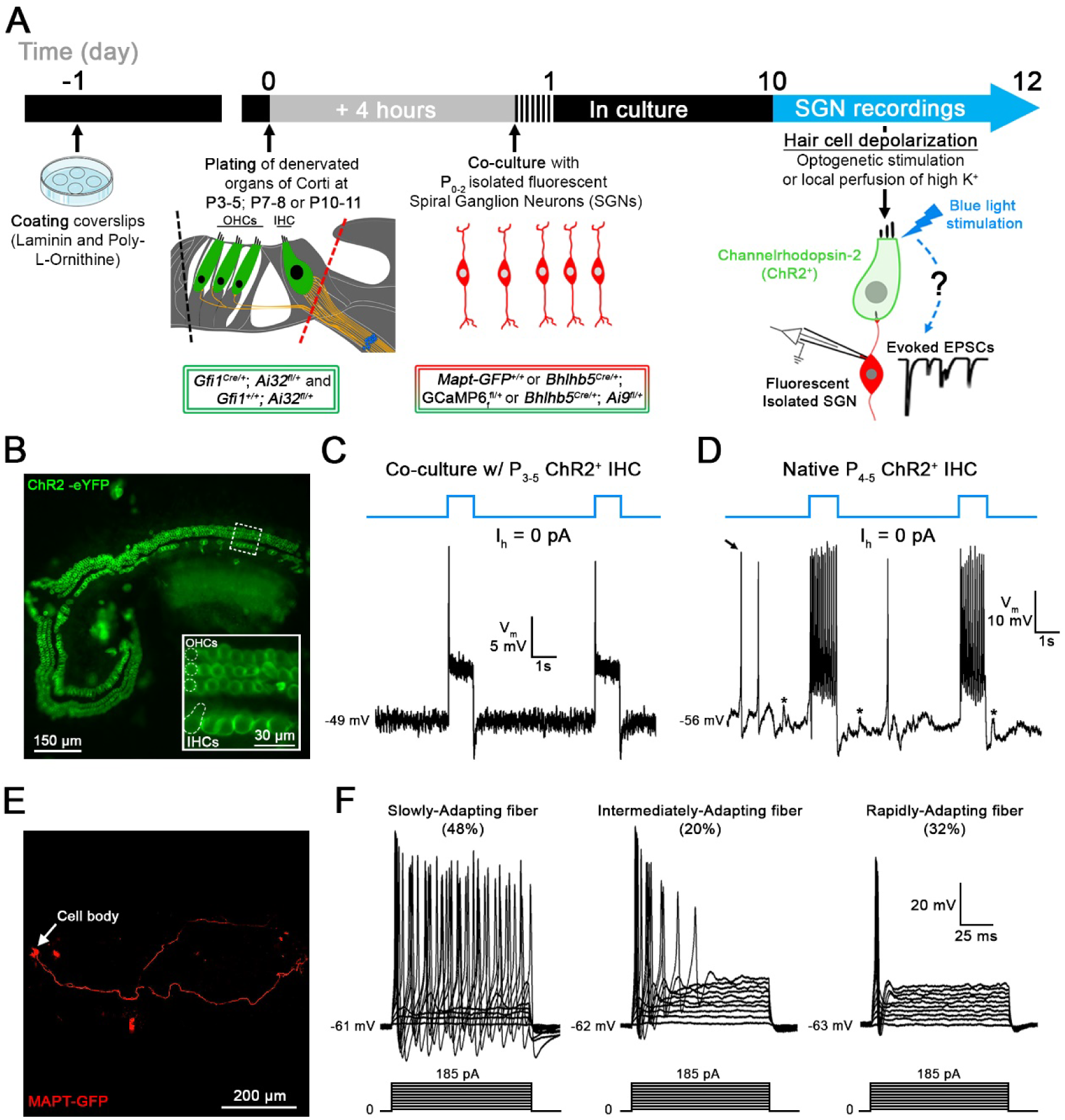
Co-cultures of denervated organs of Corti and isolated spiral ganglion neurons (SGNs) for testing regenerated hair cell synaptic function. **A**, Organs of Corti were dissected from *Gfi1^Cre/+^; Ai32^fl/+^*mice expressing Channelrhodopsin-2 (ChR2) in both inner hair cells (IHCs) and outer hair cells (OHCs) for light stimulation and from *Gfi1^+/+^; Ai32^fl/+^*mice, where hair cell stimulation was performed by the local perfusion of 40 mM K^+^. Organs of Corti were separated from the lateral wall (black dotted line) and denervated by cutting through SGN endings close to the IHCs, (red dotted line). Denervated organs of Corti were plated at 3 ages, postnatal days (P)3-5, P7-8 or P10-11. Fluorescent SGNs were isolated at P0-2 from one of three mouse lines used indiscriminately (*Mapt-GFP^+/+^*; *Bhlhb5^Cre/+^; GCaMP6f^fl/+^*or *Bhlhb5^Cre/+^; Ai9^fl/+^)*, and co-cultured with denervated organs of Corti. At 10 to 12 days *in vitro* (DIV), recordings were performed from SGN somata while stimulating hair cells. Regenerated functional synapses were identified by excitatory postsynaptic currents (EPSCs) in SGNs in response to IHC stimulation. **B**, Confocal image of a denervated P_3-5_ *Gfi1^Cre/+^; Ai32^fl/+^* organ of Corti. Inset: ChR2 is observed in IHC and OHC membranes via eYFP tag co-expressed with ChR2. **C**, In current clamp mode, I_holding_ = 0 pA, the IHC membrane potential is recorded in response to two 1s blue light pulses separated by a 4s interval (blue line). Resting membrane potential indicated at the trace. The IHC response includes an initial peak, followed by a steady-state depolarization. **D**, Same protocol as in C. IHC recording in a native P_4-5_ *Gfi1^Cre/+^; Ai32^fl/+^* acutely isolated organ of Corti. Light stimulation induces IHC depolarization and superimposed Ca^2+^ APs which are not found in cultured IHCs like in C. EPSPs (asterisks) and Ca^2+^ APs (black arrow) also occur spontaneously. **E**, Confocal image of a cultured MAPT-GFP positive SGN with soma and projecting fiber. **F**, SGNs with diverse electrical response properties in culture; ‘Slowly-Adapting’: APs slightly decrease in size throughout the pulse; ‘Intermediately-Adapting’: multiple APs only at the beginning of the pulse; ‘Rapidly-Adapting’: single AP at the pulse onset. Current clamp recordings of SGNs; 100ms long current step protocols from resting membrane potential as indicated at trace, with initial 5mV step and subsequent 10mV increasing steps.

To harvest isolated SGNs, several mouse models with fluorescently labeled SGNs were used indiscriminately: *Mapt-GFP* ^+/+^; *Bhlhb5^Cre/+^*; *GCaMP6_f_^fl/+^* (includes GFP) and *Bhlhb5^Cre/+^*; *Ai9^fl/+^* (td Tomato). Recording from such fluorescently labeled SGN somata (**Fig. 1E**) while stimulating hair cell release will assure that recorded synaptic activity originates only from newly formed hair cell/SGN synapses. SGNs were dissociated at P0-2 and added 4 h after the denervated organ of Corti had been plated. Co-cultures were kept for 10-12 days *in vitro* (DIV) before SGN recordings during hair cell stimulation were performed.

### Optogenetic stimulation depolarizes ChR2^+^ IHCs in culture

Denervated P_3-5_ organs of Corti were cultured and tight-seal, whole-cell recordings were performed from ChR2^+^ IHCs at DIV10-12, to test the approach for stimulating IHCs (**Fig. 1C, D**). The IHC membrane potential was −50 mV (−50.2 ± 4.4 mV; n = 23). One-second-long blue light pulses triggered IHC membrane depolarizations with an initial peak followed by an adapted steady state (**Fig. 1C**). The peak response occurred within 4.6 ± 0.7 ms, and ChR2^+^ IHCs were depolarized to −26.4 ± 3.8 mV (n = 22). After the initial peak, a steady state value of −42.3 ± 2.3 mV was reached with a decay time constant of 7.8 ± 2.4 ms. No response was triggered in ChR2^-^ IHCs (n = 12; data not shown), assuring that depolarization of ChR2^+^ IHCs was due to ChR2 activation.

For comparison, recordings were also performed in IHCs of acutely isolated organs of Corti whole mount preparations at P4-5 that had preserved ‘native’ IHC synapses and included the spiral ganglion (**Fig. 1D**). Here, the IHC membrane potential was about 15 mV more hyperpolarized (−64.6 ± 5.6 mV; n = 11; p <0.0001, unpaired-t test). ChR2^+^ IHCs were depolarized to a peak level of −47.4 ± 2.8 mV and a steady state level of −52.8 ± 3.2 mV (n = 4). In 6 of 10 IHCs, light-induced depolarization triggered calcium action potentials (Ca^2+^-APs) (**Fig. 1D**) and 4 of these IHCs also displayed spontaneous Ca^2+^-APs between light pulses (**Fig. 1D**, black arrow). Ca^2+^-APs have been shown to occur in immature IHCs, before the onset of hearing^36^ and are thought to trigger bursting activity in immature auditory nerve fibers, an important mechanism for refining circuitry during development of the auditory pathway^37,38^. Such Ca^2+^-APs were not observed in IHC recordings from P3-5 denervated organs of Corti in culture, even when the IHC membrane potential was hyperpolarized by current injection, to remove sodium channel inactivation (n = 17; data not shown).

In summary, although some differences regarding IHC membrane potential and evoked activity pattern were found between cultured and acutely isolated organs of Corti, the results here confirm that optogenetic stimulations of ChR2^+^ IHCs depolarize the IHC membrane potential substantially, to values that are known to induce transmitter release^39,40^.

### SGNs retain diverse electrical response properties in culture

Electrical response properties of P0-2 isolated SGNs, co-cultured with denervated organs of Corti were tested with SGN soma recordings at DIV10-12 (n = 244), and results gained from three different mouse lines were pooled. SGN somata and fibers were visualized by their fluorescence (**Fig. 1E**). Note that most SGNs tested were unlikely to be connected to IHCs, as described below. The resting membrane potential of cultured SGNs was −64 mV (−64.20 ± 4.58 mV; n = 148). The current-voltage relations typically showed robust TTX-sensitive Na^+^ inward currents (240/244) and delayed rectifier K^+^ outward currents (244/244) (data not shown). In response to a series of 100ms long current injection steps, SGN firing displayed different degrees of adaptation during the current pulse, as described before^41,42,43^ (**Fig 1F**). Accordingly, response patterns were grouped into Slowly-Adapting (48%; 78/162) (APs change in size but persist throughout pulse), Intermediately-Adapting (20%; 33/162) (multiple APs at beginning of pulse) and Rapidly-Adapting SGNs (32%; 51/162) (one AP at beginning of pulse). Slowly-Adapting SGNs displayed more negative AP thresholds compared to Intermediately- and Rapidly-Adapting SGNs (−46.87 ± 5.11 mV, n = 75 vs −42.96 ± 5.80 mV, n = 33 vs −43.14 ± 4.92 mV, n = 54; respectively, p < 0.05, Kruskal-Wallis). These different SGN response patterns could represent different maturation states of the SGNs^41^. They could also reflect different functional subtypes of SGNs which here persist after DIV10-12. Markowitz and Kalluri (2020)^43^ have found a correlation between SGN response pattern and location of SGN contact at the IHC’s synaptic pole (pillar versus modiolar), and former studies have connected these sites of contact with different physiological^44^ as well as molecular SGN properties^45;46;47^.

### New contacts between SGNs and hair cells appear in co-culture and express AMPA receptors

Earlier work has used immunolabeling of pre- and postsynaptic markers, to show that new synaptic contacts between hair cells and SGNs form in culture^10–11^. For the approach here, culture conditions were modified and two different mouse models instead of one were used for co-culture. Therefore, it was reexamined if new IHC/SGN contacts occur in co-culture.

Co-cultures of denervated organs of Corti (P3-5) and isolated SGNs (P0-2) were visualized at DIV10-12 (**Fig. 2A-C**). At that time point, a single row of IHCs and three rows of OHCs could often still be identified by shape and location of the fluorescing hair cells (ChR2-eYFP). From the randomly positioned SGN somata (arrowheads), fibers (in red) extended towards the hair cells and were seen to come into close contact with them. Typically, a single fiber made multiple contacts by branching, either with the same or with several hair cells (**Fig. 2A, B**). Often, fibers grew along multiple hair cells, appearing to be making ‘en passant’ contacts (**Fig. 2C**). This connectivity pattern is reminiscent of the developmental innervation pattern, where branched fiber endings, contacting the same or multiple hair cells are found at the beginning of the first postnatal week (P0-1) and then start to be refined between P4 and P8 to be fully pruned around the onset of hearing (∼P12)^48^.

**Figure 2.**
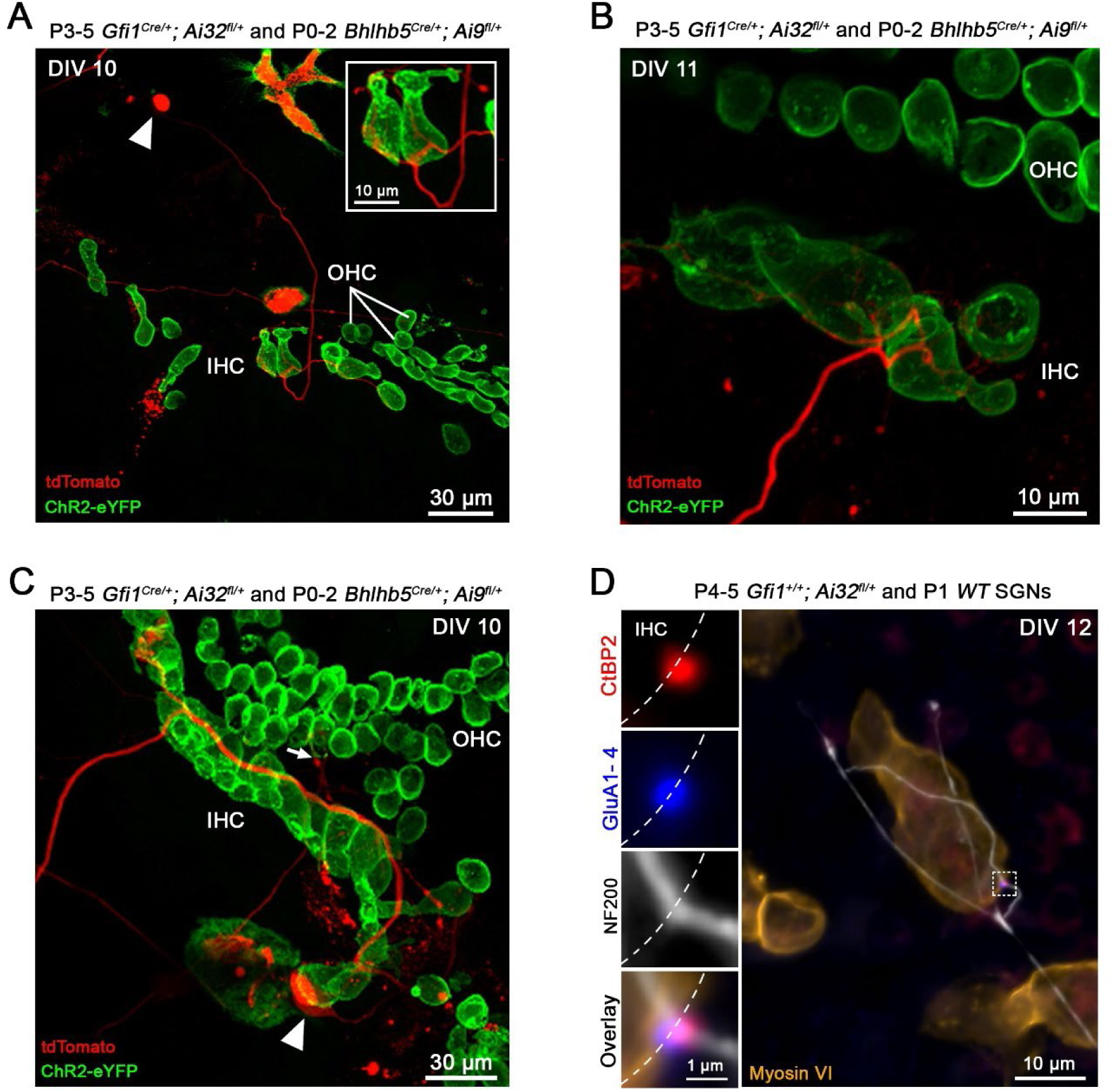
New contacts between hair cells and spiral ganglion neurons appear in co-culture and express AMPA receptors. **A-C**, Representative examples of live co-cultures at DIV10-11 showing *Bhlhb5^Cre/+^; Ai9^fl/+^*, td-Tomato expressing SGN endings (red) close to ChR2^+^ hair cells (green). In **A** and **B**, several IHCs are contacted by multiple endings from a single SGN. Arrowheads in **A** and **C** point to a SGN soma. In **C**, SGN fiber travels along the row of IHCs. Arrow points to a SGN projection reaching the OHC region. **D,** Confocal maximum-intensity projection image from an immuno-labeled co-culture at DIV 12. The insets show individual labels and overlay at a single synapse, with presynaptic ribbon (Anti-CtBP2; red) juxtaposed with postsynaptic AMPA receptors (Anti-PAN GluA1-4; blue). Anti-Myosin VI (orange) labels hair cells and Anti-NF200 (white) labels nerve fibers.

Sound signal encoding at hair cell ribbon synapses requires the close association between presynaptic ribbons, where glutamate-filled synaptic vesicles are clustered for release, and postsynaptic AMPA-type glutamate receptors (AMPA-Rs)^12;49–50^. Up until now, hair cell synapses in culture have been identified via the juxtaposition of immunolabeling for presynaptic ribbons (CtBP2, a component of the ribbon protein ribeye) and postsynaptic density markers (like PSD95)^7,10, 11^. AMPA-R labeling in young cultured cochlear tissue has been notoriously difficult and was successful here in a small set of experiments. Co-cultures of denervated organs of Corti (P3-5) and isolated SGNs (P0-2) were immunolabeled at DIV10-12. From 6 different cochlear tissues (3 mice), 26 new synaptic contacts between 18 IHCs and SGN fibers were identified by juxtaposition of CtBP2 and AMPA-R labeling (GluA1-4), providing an average of ∼1.5 synaptic contacts per innervated IHC. In such an example (**Fig. 2D**), an afferent fiber labeled with anti-Neurofilament 200 (NF200, white) is seen to meander along a row of IHCs (anti-myosin VI labeling, brown), and making a contact (marked by the dashed square) with the shown IHC. The inset magnifies the marked region of interest and shows the close apposition of CtBP2 and AMPA-Rs in a regenerated synapse. Within the set of immunolabeled tissue samples here, no OHC synapses were identified. However, during live tissue imaging, in a few instances fibers were found in close apposition with OHCs (**Fig. 2C**, arrow). Newly formed OHC synapses have been shown before by immunolabeling, but not after the first week in culture^11^.

### Hair cell stimulation activates glutamatergic synaptic currents with various waveforms in regenerated synapses

To test if regenerated synaptic contacts are functional, soma recordings were performed from SGNs in co-culture with the denervated organ of Corti (**Fig 3B**). For comparison, SGN soma recordings were performed at P4-5 in acutely isolated whole-mount preparations that included ‘native’ IHC afferent synapses (**Fig. 3A)**. Hair cell exocytosis was triggered either by one-second-long pulses of blue light stimulation or by a local perfusion of extracellular solution with elevated K^+^ (40 mM) in preparations that did not express Channelrhodopsin in hair cells^25^. Both stimulation methods provided comparable results that were pooled.

**Figure 3.**
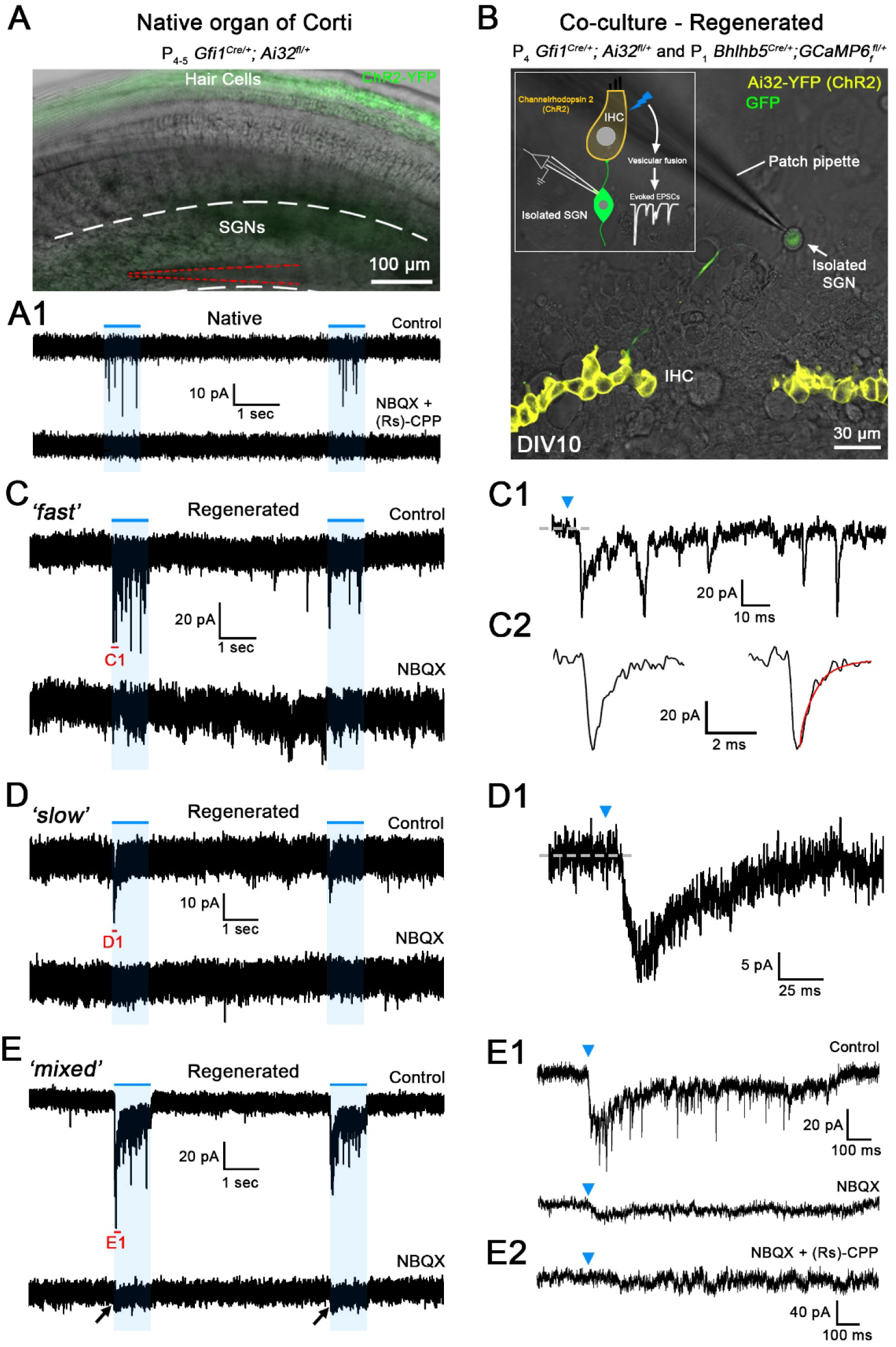
Hair cell stimulation activates glutamatergic synaptic currents with various waveforms in regenerated synapses. **A**, Superimposed difference interference contrast (DIC) and ChR2-YFP confocal image of a native acutely excised P_4-5_ *Gfi1^Cre/+^; Ai32^fl/+^* organ of Corti. ChR2^+^ hair cells in green. SG region with SGN somata is located between white dashed lines; recording patch pipette is highlighted by red dashed lines. **A1**, Trace of the SGN recording shown in **A** from a native P_4-5_ synapse, in response to two 1s long light pulses (blue lines). Holding potential −79 mV. A flurry of ‘fast’ EPSCs is observed in response to each light pulse (top trace, Control) that are blocked by the combined perfusion of NBQX and (Rs)-CPP (20 µM each), a AMPA/kainate and NMDA receptor blocker, respectively (bottom trace). **B,** A superimposed DIC and ChR2-YFP confocal image of a live co-culture at DIV 10. Row of ChR2^+^ IHCs in yellow. In this example, SGNs express the Ca^2+^ indicator GCaMP6_f_ (green) under the control of the BHLHB5 Cre promoter. Inset shows a drawing of the experimental setting. **C-E,** Trace examples of SGN recordings of regenerated synapses in co-culture using P_3-5_ denervated organs of Corti, in response to two 1s long light pulses (blue lines). Holding potential −79 mV. **C1-E1** show extended traces of C-E; their extent indicated in red **in C-E**. Blue arrowheads in C1-E1 indicate the beginning of the light pulse. Three response types with different waveforms were found: **C,** ‘fast’; **D,** ‘slow’ and **E,** ‘mixed’, the last having both fast and slow responses. Inward currents of all response types were blocked by glutamate receptor blockers; here in the examples **C1** and **D1** completely by NBQX (20 µM), in **E1** mostly by NBQX, and in **E2** completely by NBQX and (Rs)-CPP (20 µM).

In SGN recordings with native P4-5 synapses, stimulation of hair cell exocytosis triggered a flurry of exclusively ‘fast’ synaptic events (n = 7) with 10-90% rise times, median value of 1.21 ms, and 90-10% decay times, median value of 1.81 ms (149 EPSCs from 7 cells; **Fig. 4C-D)**. These EPSCs were blocked with the AMPA/Kainate-R blocker NBQX (20 mM) combined with the NMDA-R blocker (Rs)-CPP (20 mM) (n = 3) (**Fig. 3A1**), consistent with earlier recordings of AMPA/NMDA-R mediated synaptic currents in 5-7 day-old rat SGNs^51^.

**Figure 4.**
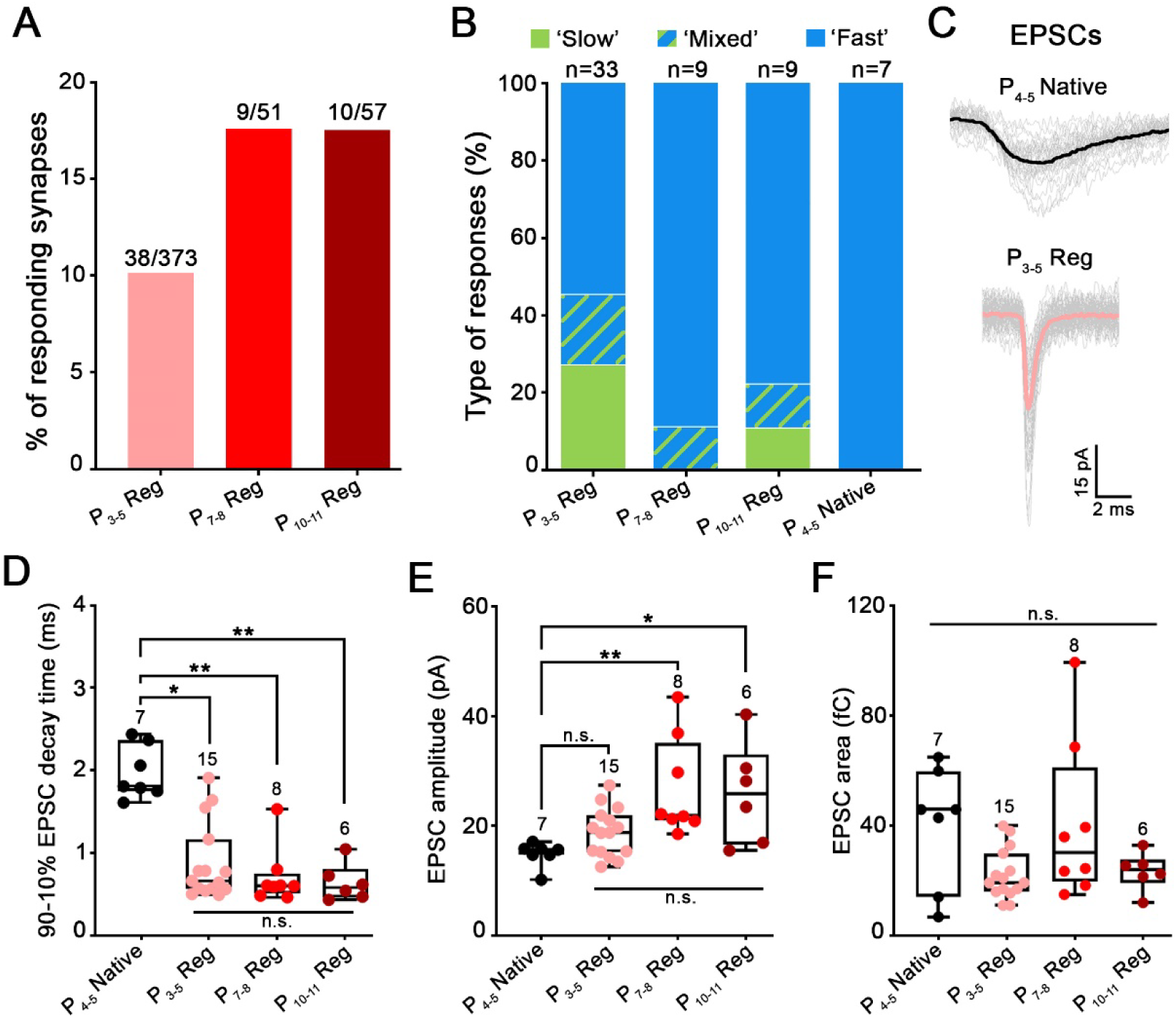
Age of the plated organs of Corti affects the maturation state of regenerated synapses. **A-F**, Properties of SGN recordings are reported for regenerated synapses in co-cultures of SGNs and denervated organs of Corti, plated at ages P3-5, P7-8 or P10-11; and for native acutely excised P_4-5_ organs of Corti (P_4-5_ Native). **A**, Percentage of SGN recordings showing regenerated postsynaptic activity in response to hair cell stimulation is higher in P_7-8_ or P_10-11_ versus P_3-5_ cultures. **B,** Distribution of ‘fast’, ‘slow’ and ‘mixed’ responses (as shown in Fig. 3) of regenerated synapses in comparison to P_4-5_ Native synapses. Contribution of ‘fast responses is higher in regenerated synapses of P_7-8_ or P_10-11_ versus P_3-5_ cultures, and 100% in P_4-5_ Native synapses. **C,** EPSC waveform of a P_4-5_ Native synapse (black) is slower that EPSC waveform from a P_3-5_ regenerated synapse (pink; P_3-5_ Reg). Individual waveforms are shown in grey and averaged in black or pink. Holding potential −79 mV. **D-F**, Comparison of EPSC waveform parameters for regenerated and native synapses. Each data point represents the median value from an individual recording. Numbers above each box indicated the number of SGN recordings used for analysis. Boxes represent the median (horizontal line), 10th and 90^th^ percentile. Whiskers represent maximum and minimum values of the distribution. *p<0.05, **p<0.01, n.s.: not significant, Kruskal–Wallis with Dunn’s post hoc test. **D**, 90-10% median decay time of native EPSCs (EPSCs^(P4–5^ ^Native)^; 1.81ms; n=7) were slower compared to regenerated EPSCs of all age conditions: EPSCs^(P3–5^ ^Reg)^ (0.66ms; n=15; p<0.05); EPSCs^(P7–8^ ^Reg)^ (0.60ms; n=8; p<0.01) and EPSCs^(P10–11^ ^Reg)^ (0.58ms, n=6; p<0.01). However, decay time was not statistically different between regenerated EPSCs of all age conditions. **E,** The median amplitude of native EPSCs (EPSCs^(P4–5^ ^Native)^: 15.59pA; n=7) was similar compared to EPSCs^(P3–5^ ^Reg)^ (18.72pA; n=15). However, EPSCs^(P4–5^ ^Native)^ were significantly smaller when compared to EPSCs^(P7–8^ ^Reg)^ (21.89pA; n=8; p<0.01) and to EPSCs^(P10–11^ ^Reg)^ (25.77pA; n=6; p<0.05). **F,** The median area of native EPSCs (EPSCs^(P4–5^ ^Native)^; 46.00fC; n=7) was similar compared to regenerated EPSCs at all age conditions: EPSCs^(P3–5^ ^Reg)^ (19.27fC; n=15), EPSCs^(P7–8^ ^Reg)^ (30.25fC; n=8) and EPSCs^(P10–11^ ^Reg)^ (24.00fC; n=6). Regenerated EPSCs also displayed similar area values at all age conditions.

When denervated organs of Corti were co-cultured with P_0-2_ SGNs, after DIV10-12, 10% of the recordings (38 of 374 SGNs) showed a synaptic response to hair cell depolarization **(Fig. 3C-E)**. Synaptic responses were divided into ‘fast’, ‘slow’ and ‘mixed’ (slow and fast combined) groups, based on their EPSC waveforms (**Fig. 3C-E**). 51% (18/33) of recordings consisted of a flurry of ‘fast’ EPSCs, with 10-90% rise times of 0.42 ms (median value) and 90-10% decay times of 0.66 ms (3346 EPSCs analyzed from 15 SGNs; **Fig. 4B-D**). These regenerated EPSCs were significantly faster than the native EPSCs (10-90% rise times: p<0.001; one-way ANOVA with Tukey’s post hoc test and 90-10% decay times: p<0.05; Kruskal–Wallis with Dunn’s post hoc test **Fig. 4D**). 25% (9/33) of recordings had a ‘slow’ response, with a single peak and slow decay occurring during the stimulus (**Fig. 3D**). 10-90% rise time median values were 16.56 ms and 90-10% decay time median values were 55.93 ms (71 EPSCs from 6 cells). 17% of recordings (6/33) showed ‘mixed ‘responses displaying a slow response superimposed by fast EPSCs (**Fig. 3E**). The time courses of slow and fast components in the mixed responses resembled those occurring in the exclusively slow or fast responses, (median values for slow component: 10-90% rise times of 19.99 ms and decay time constants of 67.32 ms (34 EPSCs analyzed); median values for fast component: 10-90% rise times of 0.47 ms and 90-10% decay times of 0.44 ms (407 EPSCs analyzed from 4 cells). Mixed responses were interpreted to be due to multiple synapses with diverse properties feeding into an individual SGN, based on the findings that afferent endings could make multiple contacts with IHCs (**Fig. 2A-C**).

Fast EPSCs in both fast and mixed responses were mediated by AMPA-Rs, as shown by complete block with the specific AMPA-R blocker CP465-022 (1 µM) (n = 6 for ‘fast’; n = 2 for ‘mixed’) and supported by complete block with the AMPA/Kainate-R blocker NBQX in additional cells (20 µM) (n = 5 for ‘fast’; n = 2 for ‘mixed’; **Fig. 3C, E**). Slow responses and slow components of mixed responses were partially or completely blocked by combinations of glutamate receptor blockers in 7 of 8 recordings (**Fig. 3D, E**). CP465-022 (1 µM), NBQX (20 µM) and (R_s_)-CPP (20 µM; NMDA-R blocker), were used in different sequential combinations, and partial block or lack of block indicated, that AMPA, kainate and NMDA receptors might all be participating in mediating the slow component to different extents, depending on the individual recording. For example, the slow component of the mixed response in **Fig. 3E** is partially blocked by the AMPA/kainate-R blocker NBQX, and completely blocked by adding the NMDA-R blocker (R_s_)-CPP. In a different example, the slow component was left unaffected by CP465-022 but completely abolished by NBQX, suggesting the involvement of Kainate-Rs in the slow component. The exact contribution of glutamate receptor subtypes to the slow component was not further investigated; further analysis of this study focused on the waveforms of fast AMPA-mediated EPSCs only (**Figs. 4 and 5**).

### Hair cell age affects the maturation state of regenerated synapses

IHC synaptic transmission is highly specialized, and single ribbon synapses show a wide range of EPSC amplitudes with some unusually large EPSC amplitudes and complex EPSC waveforms with single or multiple peaks (mono- and multiphasic)^24,25^, which are believed to occur based on more or less coordinate release of multiple release events. On the other hand, OHC synapses, like many AMPA receptor mediated CNS synapses, exhibit smaller single-peaked EPSCs^52^.

Although the IHC’s specialized release mechanism is not completely understood, it is believed to play a crucial role in proper encoding of the sound signal^24,25,26, 34, 53^. Ultimately, regenerated IHC synapses should mimic the native IHC synapse’s specialized performance. It is therefore desirable to recreate more mature and possibly IHC-type synaptic transmission in culture as a working model for further improvement of regenerative conditions.

Here we tested the hypothesis that the maturation stage of the IHC at the time of plating may affect the maturity and features of new synapses formed in culture. For this purpose, denervated organs of Corti plated at P3-5 for culture were compared to others plated at P7-8 or P10-11. Besides the age of the organ of Corti explant, experimental conditions were the same as used before, including the age of the cultured SGNs (P0-2). Interestingly, when culturing more mature denervated organs of Corti, the percentage of recorded SGNs that displayed synaptic currents in response to hair cell stimulation, increased from 10% (38 of 373; P_3-5_Reg) to 17.6% (9 of 51; P_7-8_Reg) and 17.5% (10 of 57; P_10-11_Reg) **(Fig. 4A)**. Secondly, the percentage of synapses showing solely fast EPSCs, as 100% in P_4-5_ native synapses do, increased from 48 % with P_3-5_Reg to 89 % with P_7-8_Reg and 78 % with P_10-11_Reg (**Fig. 4B**).

To compare EPSC waveforms in different experimental conditions (**Figs. 4** and **5)**, only recordings with fast EPSCs were analyzed for EPSC 90-10% decay times, amplitudes and areas (area representing the charge of the EPSC) (**Figs 4D-F)**. None of these EPSC waveform properties differed for regenerated synapses cultured with organs of Corti of different ages. However, median 90-10% EPSC decay times of regenerated synapses for all three age conditions were about 2.5 times faster compared to native EPSCs recorded from P4-5 SGNs (P_4-5_ Native:1.81 ms (n = 7; 149 EPSCs); EPSCs^(P3–5^ ^Reg)^: 0.66 ms (n = 15; 3346 EPSCs); EPSCs^(P7–8^ ^Reg)^: 0.60 ms (n = 8; 7243 EPSCs); EPSCs^(P10–11^ ^Reg)^: 0.58 ms (n = 6; 8535 EPSCs), p > 0.99, Kruskal–Wallis with Dunn’s post hoc test; **Fig. 4D**). This difference is illustrated with example EPSC waveforms from a native P_4-5_ synapse (P_4-5_ Native) and from a regenerated synapse with a P_3-5_ denervated organ of Corti (P_3-5_ Reg; EPSCs^(P^^3–5^ ^Reg)^;**Fig. 4C**). The presence of faster EPSCs in regenerated synapses might indicate some form of maturation in culture, that could occur by a reduction in the NMDA component as found in early postnatal development^51^ and/or a change in AMPA receptor subtype expression, as AMPA receptor mediated EPSCs in SGNs speed up during development^25^. Similarly, amplitudes of EPSC^(P7–8^ ^Reg)^ and EPSC^(P10–11^ ^Reg)^ were significantly larger compared to native EPSCs recorded from P4-5 SGNs (P_4-5_ Native; respectively, p < 0.01 and p < 0.05, Kruskal–Wallis with Dunn’s post hoc test), whereas EPSC^(P3–5^ ^Reg)^ were not (p = 0.88, Kruskal–Wallis with Dunn’s post hoc test, **Fig. 4E**).

In summary, when older organs of Corti were plated, more synapses showed the more mature ‘fast’ responses. However, the basic measures of EPSC waveform like amplitude, area, and decay time, were not obviously affected. To further probe for possible differences, a more detailed analysis of EPSC waveforms was performed next.

### EPSC waveforms at synapses regenerated with older IHCs reveal properties closer to mature IHC ribbon synapses

Deconvolution analysis of EPSC waveforms was performed^34^, probing for possible changes in the mechanism of release that may occur when synapses mature. Only EPSCs of ‘fast’ responses were analyzed. The approach is illustrated in **Fig. 5A1**. This method relies on the averaging of monophasic EPSCs (with a smooth rise, a single peak and exponential decay) to compute a’ kernel’ that is presumed to reflect a single release event (**Fig. 5A2**). ‘Multiphasic’ EPSCs with more complex waveforms, are assumed to be the linear sum of multiple asynchronously-released kernels, representing a varying sequence of neurotransmitter release events (**Fig. 5A3-A5**). The sum of kernels required to fit an EPSC is computed (orange dotted line), using the lasso optimization method^35^. This analysis allows estimation of the timing, number and relative sizes of presumed release events making up an EPSC, as illustrated by the dotted peaks of the green lines in **Fig. 5A**. **Fig. 5A** illustrates that individual recordings can include EPSCs with varying numbers of events (kernels) per EPSC, as indicated by the numbers above the EPSCs.

**Figure 5.**
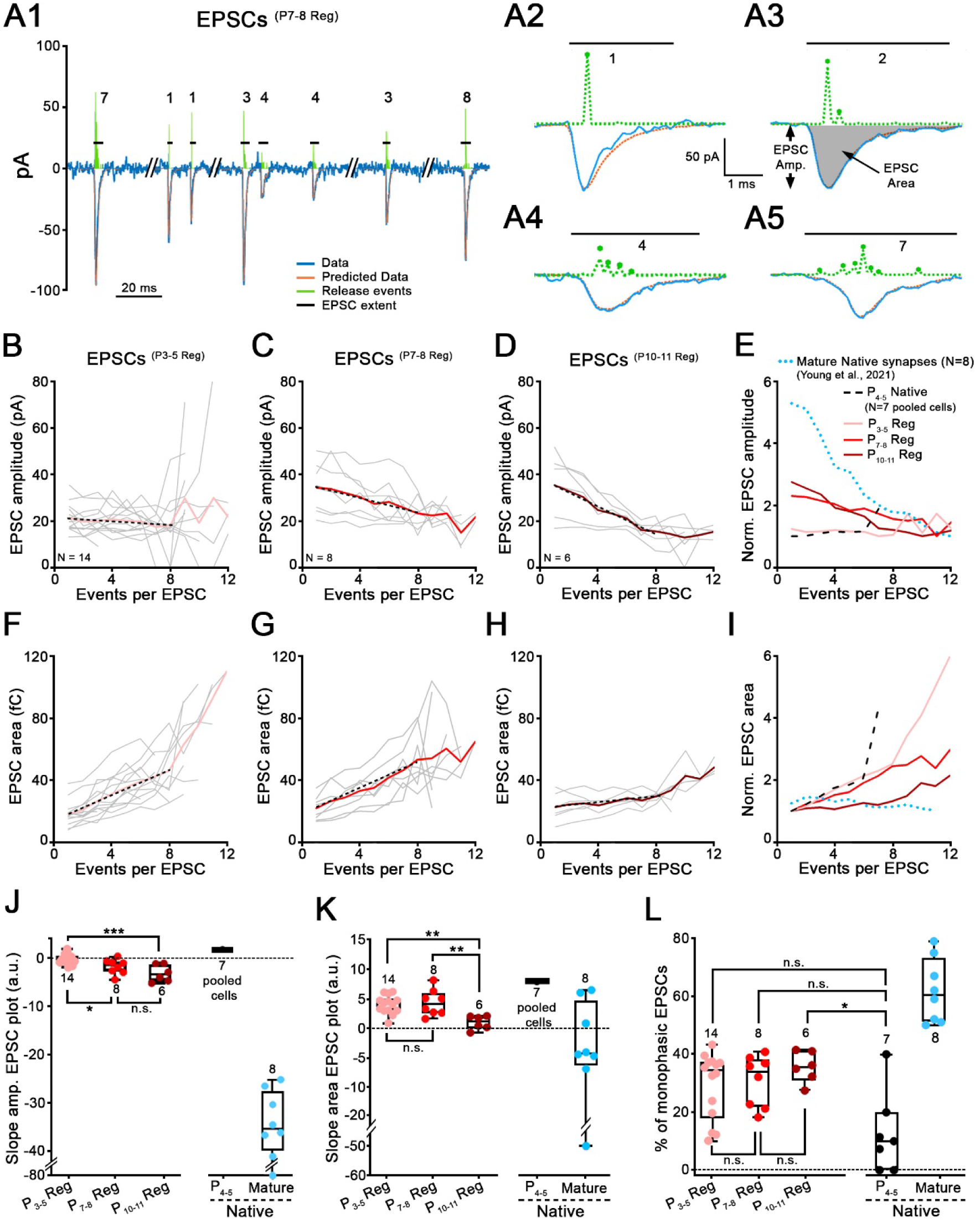
EPSC waveforms at synapses regenerated with older IHCs reveal properties closer to mature IHC ribbon synapses. **A**, Modeling of EPSC waveforms using deconvolution. **A1,** Trace (blue line) of a SGN recording with 8 exemplar EPSCs^(P7–8^ ^Reg)^ recorded from a regenerated synapse. Holding potential: –79 mV. P_7-8_ denervated organ of Corti was plated for this co-culture with P_0-2_ SGNs. Four EPSCs from this recording are shown on extended time scales in **A2-A5** (blue lines). **A2,** Monophasic EPSCs like this example, were averaged to create a kernel (standardized release event) for individual recordings. Kernels (amplitude and time-of-occurrence) are depicted in green dashed line above each EPSC. The fits calculated from the event sequences are shown in orange. EPSC amplitude (EPSC Amp.) and EPSC area (grey filled area) are defined in **A3**. Horizontal black lines represent EPSC extent and numbers indicate the smallest number of events (i.e., kernels) that best fit this EPSC. **B-D,** Mean values of regenerated EPSC amplitude are plotted against the number of events per EPSC, for three co-culture conditions, P_3-5_ (**B,** n=14 synapses, pink), P_7-8_ (**C,** n=8 synapses, red) and P_10-11_ (**D,** n=6 synapses, purple) denervated organ of Corti. Grey thin lines represent individual recordings and bold colored lines represent the averages. Black dotted lines represent the fit of the data, including only data with 1-8 events per EPSC, providing the slope values in **J**. **E**, EPSC amplitude versus events/EPSC plots were normalized to their minimum value and superimposed for different conditions. These include average traces for regenerated synapses from **B**-**D, (**P_3-5_ Reg, P_7-8_ Reg, P_10-11_ Reg; pink, red and purple), for immature P_4-5_ Native synapses (black dashed lines) and mature ribbon synapses (blue dotted lines, data from^34^). For EPSCs^(P4–5^ ^Native)^, EPSCs from the 7 recordings were pooled. **F-I,** Same as **B-E**, but with EPSC area plotted against number of events per EPSC. **J,** Slopes of the EPSC amplitude versus the number of events per EPSC calculated from **B-E**, are shown for each condition. EPSCs^(P4–5^ ^Native)^: 1.79 (n=7 pooled cells); EPSCs^(P3–5^ ^Reg)^: −0.32 (n=14); EPSCs^(P7–8^ ^Reg)^: −1.44 (n=8); EPSCs^(P10–11^ ^Reg)^: −3.26 (n=6) and EPSCs^(mature^ ^native)^: −35.76 (n=8; data from^34^). One-way ANOVA with Tukey’s post hoc test. **K,** Slopes of the EPSC area versus the number of events per EPSC calculated from **F-I**, are shown for each condition. EPSCs^(Native^ ^P4–5)^: 8.00 (n=7 pooled cells); EPSCs^(P3–5^ ^Reg)^: 4.33 (n=14); EPSCs^(P7–8^ ^Reg)^: 4.01 (n=8); EPSCs^(P10–11^ ^Reg)^: 1.7 (n=6) and EPSCs^(mature^ ^native)^: −4.32 (n=8; data from^34^). One-way ANOVA with Tukey’s post hoc test. **L,** Percentage of monophasic EPSCs per recording is shown for each condition. EPSCs^(Native^ ^P4–5)^: 10% (n=7); EPSCs^(P3–5^ ^Reg)^: 34.35% (n=14); EPSCs^(P7–8^ ^Reg)^: 34.04% (n=8); EPSCs^(P10–11^ ^Reg)^: 35.64% (n=6) and EPSCs^(mature^ ^native)^: 60.50% (n=8; data from^34^). Kruskal–Wallis with Dunn’s post hoc test. **J-L**, Each data point represents an individual recording. Number of SGN recordings used for analysis is indicated. Boxes represent the median (horizontal line), 10th and 90^th^ percentile. Whiskers represent maximum and minimum values of the distribution. *p<0.05, ***, **p <0.01, p<0.001, “n.s.”: not significant.

It was shown before that EPSCs recorded in native more mature SGN synaptic terminals have very specialized release properties^24–26, 34^. To illustrate such features, summary data from 2– 4-week-old synapses from hearing animals are replotted from Young et al. (2021)^34^ in **Figs. 5E, I** (blue dotted lines). For this data set, the areas of EPSCs, representing EPSC charge, remained close to unchanged (only slightly decreasing) with increasing numbers of events per individual EPSC at a given synapse (**Fig. 5I**). This result implies that a fixed amount of neurotransmitter is released for each EPSC and that multiphasic EPSCs reflect the desynchronization of that transmitter release into several boluses. Thus, the amplitude of EPSCs must decrease with the increasing number of events per EPSC as shown in **Fig. 5E**, for the EPSC area to remain unchanged.

Results from acutely isolated tissue (P4-5) with immature native synapses are also shown for comparison (P_4-5_ Native; black dashed line in **Fig. 5E, I**); here all recordings were pooled (n = 7; 149 EPSCs), due to the small numbers of EPSCs per recording. The small sample size also causes an uptick in the graphs for values > 6 events per EPSC; these values were not included in the interpretation of the data. For immature native synapses, EPSC area increased with higher numbers of events per EPSC, whereas the EPSC amplitudes stayed about constant (very slightly increased). One interpretation of this result is that additional events in EPSCs may come from additional release pools, increasing area when similarly sized events are added. These data suggest a different mode of transmitter release for early postnatal synapses compared to more mature several weeks old synapses.

The deconvolution analysis of EPSCs was performed for the three experimental groups resulting in regenerated synapses (same as in **Fig. 4**), with the denervated organ of Corti plated at P3-5, P7-8 or P10-11. The dependence of EPSC amplitude and EPSC area on number of events/EPSC was plotted for individual recordings (in gray) and averages across recordings (in color) (**Figs. 5B-D; 5F-H**). The averages of every condition were replotted in summary panels, for comparison with data from native immature and more mature synapses (**Fig. 5E, I)**. The slopes of the average traces were calculated by a linear fit to the corresponding data over the range of event numbers 1 to 8 only (illustrated by dotted lines in **Figs. 5B-D, F-H**), not using the noisier data for larger event numbers. Statistical tests were performed to compare P3-5, P7-8 or P10-11, and immature and more mature native synapse results were plotted for qualitative comparison in **Fig. 5**, but not included in the statistical analysis of plotted slopes, as data of native immature synapses had to be pooled providing a single value and as experimental conditions had been somewhat different for native mature compared to regenerated synapses.

Interestingly, properties of regenerated synapses created with plating more mature organs of Corti showed a clear trend for both EPSC amplitude and area data in the direction of those from hearing animals with more mature ribbon synapses (**Fig. 5J, K**). With increasing number of events/EPSC, EPSCs^(P3–5^ ^Reg)^ amplitudes stayed close to constant, like EPSCs^(P4–5^ ^Native)^ amplitudes, (**Fig. 5B, E**; light pink line and black dashed line). EPSCs^(P7–8^ ^Reg)^ and EPSCs^(P10–11^ ^Reg)^ amplitudes decreased with increasing number of evens/EPSC (**Fig. 5C-E**, red and purple lines), and slopes were significantly more negative for both conditions compared to EPSCs^(P3–5^ ^Reg)^ (p < 0.05 and p < 0.001 respectively, one-way ANOVA with Tukey’s post hoc test, **Fig. 5J**), trending towards the behavior of more mature native synapses (**Fig. 5E**, blue dotted line).

Secondly, with increasing number of events/EPSC, EPSC area increased in the regenerated synapses at all three ages (**Fig. 5F-5H**). EPSCs^(P3–5^ ^Reg)^ and EPSCs^(P7–8^ ^Reg)^ area data qualitatively resembled more closely those of EPSCs^(P4–5^ ^Native)^ and had significantly steeper slopes than EPSCs^(P10–11^ ^Reg)^ (**Fig. 5I**; **Fig. 5K**; p < 0.01, one-way ANOVA with Tukey’s post hoc test). The EPSCs^(P10–11^ ^Reg)^ area plot with the older organ of Corti plated, with a shallow slope, trended towards the behavior of more mature native synapses, where EPSC areas are close to constant with increasing numbers of events/EPSC.

It has been reported previously that the fraction of monophasic EPSCs in individual recordings increases with postnatal age, (second versus third postnatal week)^25^. However, the three experimental groups with regenerated synapses P3-5, P7-8 or P10-11 did not show significant differences in percentage of monophasic EPSCs (P_3-5_Reg: 34.35%, P_7-8_Reg: 34.04 and P_10-11_Reg: 35.64%). A weak trend was found in comparison to immature native EPSCs^(P4–5^ ^Native)^ (median: 10%), as only EPSCs^(P10–11^ ^Reg)^, with the older organ of Corti plated, showed a significantly higher percentage of monophasic EPSCs (p < 0.05, Kruskal–Wallis with Dunn’s post hoc test), again, trending towards the even higher fraction of monophasic EPSCs (median: 60.50 %) in native synapses from hearing animals^34^.

## DISCUSSION

The study here utilizes IHC optogenetic stimulation and recordings from fluorescing SGNs *in vitro*, showing that newly formed IHC synapses are indeed functional, exhibiting glutamatergic excitatory postsynaptic currents and show qualitative release properties reminiscent of native mature IHC synapses. This newly developed functional assessment of regenerated IHC synapses therefore provides a powerful tool for testing approaches and compounds to improve synapse regeneration.

### Limitations of the in vitro approach

The *in vitro* approach comes with some limitations. 1) Immature pre-hearing hair cells (P3-11) and SGNs (P0-2) were used, for best survival in culture. These choices will not completely reflect how synapse regeneration might occur in the mature cochlea. However, some important properties were still found intact, even under these conditions: At a prehearing age, IHCs in culture developed properties reminiscent of the specialized IHC release mechanism (see below). SGNs plated at an immature stage still displayed the response properties of diverse subgroups of type-I SGNs^22, 41, 43, 44^. 2) Supporting cells of the Greater Epithelial Ridge were mainly removed when denervating cochlear tissue. However, these supporting cells play important roles during cochlear development (for review^54^), by promoting SGN outgrowth and survival^55–57^, modulating intrinsic SGN properties^32, 43, 58^ and regulating postnatal cochlear spontaneous activity involved in circuitry refinement^37, 59–61^. 3) IHC Ca^2+^ APs have been shown to regulate ribbon synapse maturation^62–64^, but are absent in cultured IHCs. Like the absence of other presynaptic inputs^45, 46, 65^, this could affect synapse maturation. However, despite these limitations, regenerated synapses exhibited functional glutamatergic transmission, with qualitative properties reminiscent of native mature IHC ribbon synapses, validating the approach for gaining insights into mechanisms underlying synapse regeneration and for testing potential regenerative reagents.

### Glutamatergic transmission in regenerated synapses

Native IHC synapses are typically identified by immunolabeling of presynaptic ribbons (CtBP2) and juxtaposed postsynaptic AMPA-Rs^49, 50, 66–68^. Native IHC synaptic currents exhibit ‘fast’ kinetics on the millisecond scale, unusually large amplitudes and have been shown to be mediated by AMPA-Rs^24–26, 49, 69, 70^, and additionally by NMDA-Rs during early postnatal development^51, 71, 72^. Similarly, regenerated IHC synapses *in vitro* showed juxtaposed CtBP2/pan-AMPA-R immuno-puncta and a large fraction of EPSCs with ‘fast’ AMPA-R-type properties. EPSC waveforms with organs of Corti plated at P3-11 and recorded after DIV10-12, were slightly faster compared to native synapses recorded in whole mount preparations at P4-5, possibly due to further maturation during the time in culture that may have resulted in changed contributions and subunits of NMDARs^51^ and AMPA-Rs ^25, 73–76^.

Additional to ‘fast’ EPSCs, regenerated synapses showed ‘slow’ EPSCs with a tens of milliseconds time scale for rise and decay times, which based on pharmacology included a varying combination of AMPA-, NMDA- and Kainate-R mediated components. Such slow EPSCs appeared at a higher percentage with the youngest organs of Corti (P3-5) plated, suggesting that this phenotype represents a more immature state of regenerated synapses. Gluk1-5 Kainate-R have been found at mature IHC post-synapses by immunolabeling, and mRNA for Kainate-Rs with subunit specific developmental expression profiles have been found in SGNs^45, 47, 68, 77^. A physiological assessment of the role of and time course of Kainate-R responses at developing or mature IHC synapses, is not available at this point. NMDA-R subunit GluN1 and GluN2a are expressed by developing IHC-SGN synapses^70, 72^ ^78^. During cochlear maturation, GluN1 expression decreases and GluN2a is replaced by GluN2b, 2c and 2d (for review^78, 79^). Interestingly, after excitotoxic-trauma, *in-situ* hybridization revealed the re-expression of the GluN1 subunit in SGNs, and its blockade by pharmacology results in a delay in fiber regrowth and function recovery^80^. Recently, Lithium Chloride, known to downregulate GluN2b subunit expression, injected through the round window 1 day after noise exposure, led to the rescue of ribbon synapses in rats^81^. This indicates that regulation of NMDAR subunit expression may not only be involved during synaptogenesis but also may take part in synapse regeneration. In summary, data from the present study suggest that the expression of different glutamate receptor subtypes may occur during synapse regeneration. Further studies are needed to determine, if the appearance of NMDA-Rs and Kainate-Rs besides AMPA-Rs represents a recapitulation of synapse development, or is specific to the process of regeneration.

### Culturing older IHCs results in regenerated synapses with more mature, IHC-like release properties

When older IHCs were cultured with SGNs, a more mature synaptic phenotype was found in regenerated synapses. With older IHCs, a higher percentage of SGNs formed functional synapses, and more such synapses had solely fast EPSCs. With older IHCs, regenerated synapses had significantly larger EPSC amplitudes and a higher percentage of monophasic EPSCs compared to younger native IHC synapses. Similar changes of EPSC properties have also been observed for native ribbon synapses during development^25^.

IHC transmitter release at mature IHC synapses has been shown to operate by more or less coordinate release of a ‘unit/bolus’ of neurotransmitter, creating mono- and multiphasic EPSCs^25, 26, 34^. This feature is indicated by EPSC area staying constant with larger numbers of events per EPSC. Interestingly, this property of release is not found when creating regenerated synapses with young IHCs in culture (P3-5), suggesting that in P3-P5 IHCs the coordinate release mechanisms has not developed yet. However, it is impressive that with older IHCs, coordinate release can be recreated in regenerated synapses, even *in vitro*.

However, although regenerated synapses *in vitro* showed coordinate transmitter release, EPSC amplitudes were on average up to ∼10-times smaller compared to mature native IHC synapses, as well as were those of native your synapses at P4. One simple possible explanation for this difference may be due to the fact that EPSCs in native mature IHC synapses were recorded close to the site of release, in IHC afferent endings^24, 25^, whereas in the study here, EPSCs were recorded in SGN somata, likely a couple of hundred micrometer away from the synapse, and transmitted by an immature unmyelinated peripheral axon, as afferent ending recordings in this approach with newly formed fragile synapses are not feasible. Optogenetic IHC stimulation in comparison to IHC depolarization by 40 mM K^+^ solution evoked EPSCs with similar amplitudes, and therefore is unlikely to have caused small EPSC amplitudes. To be able to make a clear statement regarding differences in EPSC amplitude, the same type of SGN recording would be needed at different IHC ages and in native and culture conditions.

### Other synapse regeneration models

New insights from functional testing of regenerated cochlear ribbon synapses will also inform other disease models where synapse regeneration is needed for restoring function. For example, in certain pathologies of visual function, photoreceptors may degenerate or lose their ribbon synapses^82^. *In vivo* studies have demonstrated that after transplantation, photoreceptor precursor cells (PPCs) can make new synaptic contacts in a host retina, and in some instances improve visual acuity^83, 84^. Recently, the engraftment of a genetically modified retina derived from human embryonic stem cells (hES cells) restored electrical activity of host ganglion cells in nude rats, based on multi-electrode array recordings^85^. An *in vitro* organotypic retina mouse model allows for checking the functional integration of a graft into host tissue, based on the labeling of specific synaptic proteins^86^. In a cochlear *in vitro* model, neural progenitor cells have been shown to acquire new synaptic connections with hair cells using immunolabeling^12^, and the approach reported here will now allow for testing the specific properties of such newly formed synapses. Another promising regenerative strategy is to promote the trans-differentiation of inner supporting cells into cochlear hair cells^87^. In the retina, Müller glial cells could potentially re-enter the cell cycle and be transdifferentiated into neurons (For review^88^). Such newly created sensory cells and peripheral neurons will need to be tested for proper synaptic function. Promoting regeneration in the CNS has been challenging, however, some progress has been made towards nerve growth and sometimes some functional recovery^89, 90^. One example is the blocking of the Repulsive Guidance Molecule A (RGMa). Intrathecal injection of the RGMa antibody has been performed in rats following thoracic spinal cord hemi-section and axonal growth along with function recovery were observed^91^. In the present study RGMa protein was blocked to improve IHC/SGN synapse formation, as shown before^11^. Organotypic cell culture models of cerebral or spinal cord tissues have been used for investigating their 3-D architecture^92^, mimicking neurodegenerative disease models^93^, testing the functionality of a regenerated network, and most importantly, developing new therapeutical tools^94^. In summary, multiple fields are developing approaches to better inform sensory cell, nerve fiber and synapse regeneration approaches. The present work provides first insights into the functional properties of individual regenerated synapses, and therefore has a high potential to provide new insights and cross-disciplinary exchange with the general regenerative field.

## ACKNOWLEDGEMENTS

This work was supported by the National Institute on Deafness and Other Communication Disorders Grants R01DC006476 to EG, the David M. Rubenstein Fund for Hearing Research to EG, the Geraldine Dietz Fox Endowed Research Fund to EG, the Hearing Health Foundation grant to PV, R01 DC007174 to AE. We thank Dr. Mingji Tong, Eaton Peabody Laboratory, Massachusetts Eye and Ear, Boston MA, for training in cell culture models with regenerated synapses.

